# Hybrid quantum dot - collagen extracellular matrices for in situ optical monitoring of cardiomyocyte activity by two-photon fluorescence

**DOI:** 10.1101/2020.03.18.990846

**Authors:** Stijn Jooken, Yovan de Coene, Olivier Deschaume, Olga Krylychkina, Thierry Verbiest, Koen Clays, Geert Callewaert, Carmen Bartic

**Affiliations:** Soft Matter and Biophysics Unit, Department of Physics and Astronomy, KU Leuven, Celestijnenlaan 200 D, 3001 Leuven, Belgium; Department of Life Science Technologies, imec, 3001 Leuven, Belgium; Molecular Imaging and Photonics, Department of Chemistry, KU Leuven, 3001 Leuven, Belgium; Department of Cellular and Molecular Medicine, KU Leuven Campus Kulak, 8500 Kortrijk, Belgium

**Keywords:** quantum dot, collagen, two-photon fluorescence, cardiomyocyte, optical readout

## Abstract

The incorporation of functional nanoparticles in scaffolds for tissue constructs has led to the creation of artificial extracellular matrices that more accurately mimic the cues present in the native microenvironment of developing tissue. Additionally, light-sensitive inorganic nanoparticles can act as cell biosensors and report on the physiological parameters during tissue growth and organization. In this work, we functionalized collagen nanofibers with semiconductor quantum dots (QDs) and thereby created artificial extracellular matrices that can optically report on cardiomyocyte activity based on QD two-photon fluorescence. We have applied these optically-addressable nanofiber matrices to monitor activities of primary cardiomyocytes and compared the optical responses with patch-clamp data. Combining the long-term stability of QD fluorescence with the deeper light penetration depths achievable through multiphoton imaging, this approach can be used for continuous monitoring of cellular functions in cardiac tissue engineering.

**Abstract Figure:** Concept illustration: optical readout of cardiomyocyte activity with QD-functionalized collagen networks. Whole-cell current-clamp mode is used here to simultaneously monitor changes in the transmembrane voltage while the QD two-photon fluorescence is recorded.

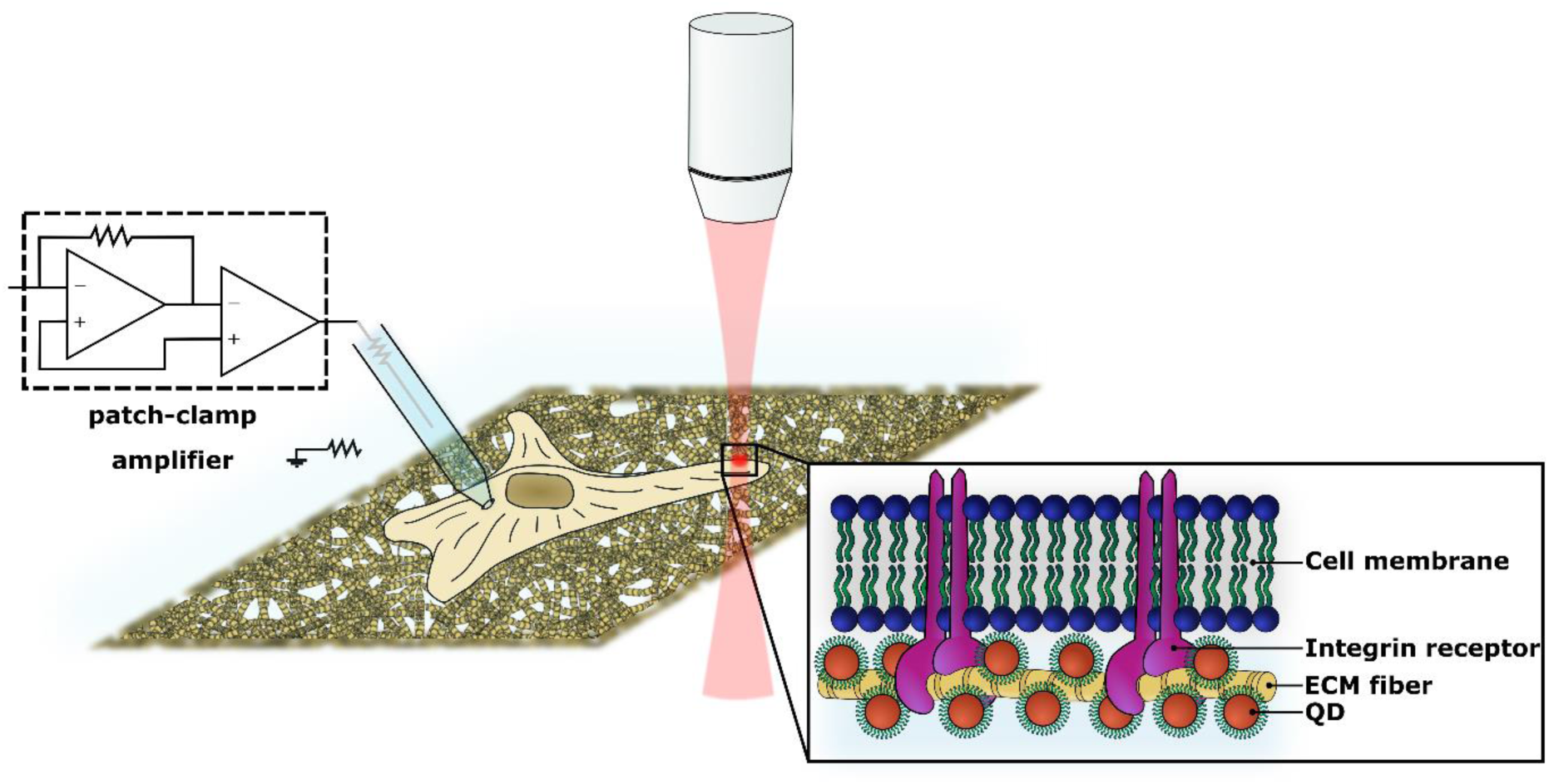

## 1. Introduction

The detection of electrical activity and contractile motion of cardiomyocytes is essential in understanding cell physiology in health and disease, performing drug screening and toxicity studies, as well as in monitoring physiological responses in tissue engineering constructs. A plethora of techniques have been proposed to assess the activity of either isolated ventricular cardiomyocytes or hiPSC (human induced pluripotent stem cell) derived cardiomyocytes, at the single cell or cluster level *in vitro*. In pharmaceutical drug development and toxicity studies, the golden standard for screening cardiomyocyte electrophysiological functionality is automated patch clamp^[1]^. Being invasive, this method only allows for single and short term measurements for each cell. Microelectrode array (MEA)^[2]^ systems enable both intra- and extracellular measurements of electrical activity, but require clusters of cardiomyocytes and are not suitable for 3D tissue preparations. Also a large variety of optical imaging techniques have been proposed for monitoring the activity of electrogenic cells, based on linear and nonlinear light-tissue interaction modalities in combination with a variety of reporter probes, some genetically encoded^[3–6]^. Still, the significant probe photobleaching accompanying the optical imaging is a major limitation for the long-term monitoring of cellular preparations.

Additionally to action potentials, the contractile properties of cardiomyocytes hold important information on the status of the tissue-engineered cardiac patch, as (drug-induced) cardiac toxicity often manifests itself not only through faulty electrical signal propagation but also affects mechanical strain, beat frequency (arrythmias) and dysregulation of calcium cycles^[1,7]^. Emerging methods to assess contractile functionality more comprehensively include optical flow-based analyses where through image processing of bright field microscopy videos^[8,9]^ or interference patterns in lens free imaging (LFI)^[10,11]^, force-vectors are generated that represent the contractile motion of the cardiomyocyte. These techniques require image post-processing and the significant computational time prevents their ability in real-time contraction monitoring. Moreover, chronotropic (altering beat rate) and inotropic (altering contraction force) effects of drugs on cardiac contraction can also be investigated using impedance spectroscopy^[12–14]^, while contraction forces and energies are often measured using flexible posts and force transducers where pre- and afterload can be manipulated^[15–17]^. All these techniques are only applicable for 2D cell monolayers. In this work we demonstrate a straightforward optical approach for monitoring cardiomyocyte contractions suitable for long-term monitoring and 3D cellular preparations using hybrid matrices combining extracellular matrix (ECM) protein nanofibers and quantum dots (QDs).

In vitro, extracellular matrices based on fibrous proteins or synthetic nanofibers can provide cells with the mechanical and biochemical cues that are present in the natural tissue and regulate cell behavior^[18]^. Moreover, in the last decade the incorporation of nanoparticles allowed to more closely mimic properties of natural ECMs^[19–22]^. Except for cue providing, the inorganic nanoparticles allow incorporating sensing modalities into extracellular matrices, not only to deliver stimuli, but also report the actual status of the electrogenic activity and contractility of the tissue-engineered cardiac patch. The monitoring function would notably impact the success of the regenerative processes, leading to better control over tissue development. The environment-sensitive optical properties of QDs make them versatile probes for locally evaluating the effects of cellular activity (e.g. pH and temperature changes, membrane voltage^[23,24]^, intracellular calcium levels^[25]^ or cytokine^[26]^ and neurotransmitter release^[27]^). Their potential to act as membrane voltage probes in electrogenic cells has been hypothesized and demonstrated in FRET- and ET-based configuration^[23,28–30]^. Compared to organic dyes and fluorescent proteins, QDs are much less affected by photobleaching, thus much more suitable for long-term experiments, exhibit sharp emission spectra – important for straightforward signal deconvolution in multiprobe imaging, and much larger absorption cross-sections for two-photon fluorescence excitation – around 50 000 GM at 900 nm for QDs^[31,32]^ versus values up to an order of magnitude of 1000 GM for dyes and proteins^[33,34]^ and organometal complexes^[35,36]^. To our knowledge, so far, no successful data on QD direct cellular activity imaging in primary mammalian cells has been reported. One major challenge is thought to be the difficulty to achieve high proximity, stable membrane localization of QDs[28,37,38].

To enhance the membrane localization, we attach QDs to fibrillar collagen, abundant in natural ECMs and essential for tissue strength and integrity^[39]^. Collagen fibers display functional domains specific to various cell membrane receptors, among others the α1β1 and α1β2 integrins^[40]^. Therefore, we hypothesized that fibrillar ECM proteins can act as biological templates for efficient QD localization in view of cell activity monitoring.

We report on the optical detection of contractile activity of rat cardiomyocytes by monitoring changes in the two-photon fluorescence of CdSe/ZnS core-shell QDs attached to collagen nanofibers and validate the data against patch clamp recordings of similar parameters in the presence of the positive chronotropic agent epinephrine.

## 2. Results and discussion

### 2.1. Hybrid collagen - quantum dot extracellular matrices

Figure 1 schematically illustrates the preparation of a hybrid QD-collagen scaffold. Octadecylamine-stabilized CdSe/ZnS core-shell QDs (Sigma Aldrich) with an emission wavelength of 630 nm (Figure S1) are solubilized in chloroform (CHCl3) and transferred to water by a polymer encapsulation step to improve biocompatibility for cell experiments (see Materials and Methods). Encapsulation into poly-styrene-co-maleic anhydride (PSMA) ensures high colloidal stability and reduces the quantum yield (QY) loss accompanying the water transfer procedure. After phase transfer, the QY of the PSMA encapsulated QDs in 50 mM HEPES buffer at pH 7.4 amounts to 18 ± 3 %. The average diameter of the nanocrystals as measured by dynamic light scattering, increases from 14.3 ± 0.2 nm to 24.2 ± 0.5 nm after PSMA encapsulation, as shown in Figure S1a.

**Figure 1.**
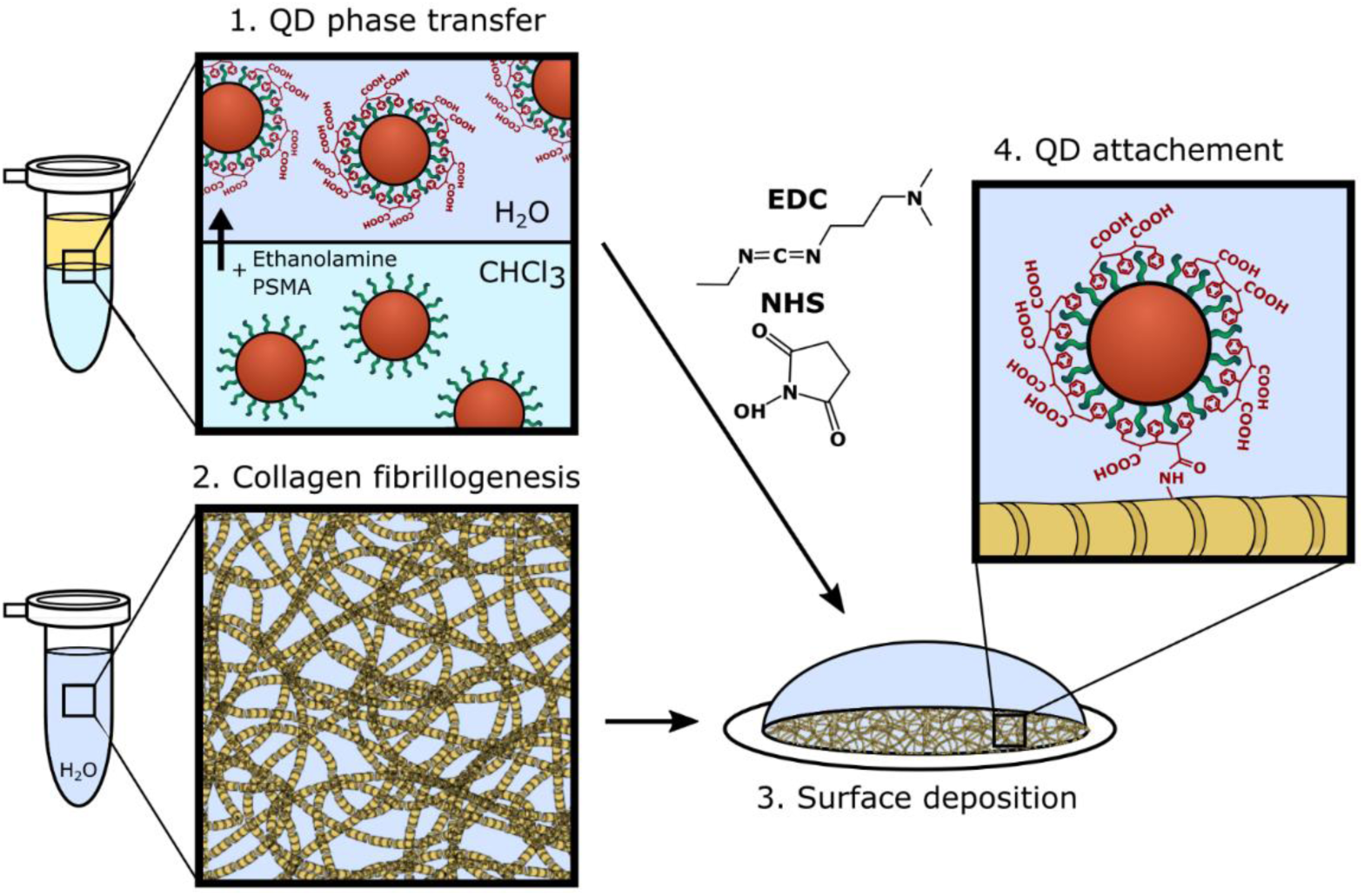
Process flow for scaffold preparation. (1) Quantum dots (QDs) are phase transferred from chloroform to water by poly-styrene-co-maleic-anhydride (PSMA) encapsulation; (2) Collagen fibers are self-assembled in phosphate buffered saline (PBS) starting from monomeric collagen; (3) The collagen network is deposited onto a poly-L-lysine (PLL) coated glass coverslip; (4) The water-soluble QDs are covalently bound to the collagen scaffold through covalent crosslinking.

Type I collagen fibers are allowed to self-assemble from bovine monomeric collagen in solution for a period of 4 hours at 37°C (see Material & Methods). The resulting collagen gels are then ultrasonicated, centrifuged for 2 min at 6000 rpm and the pellet is resuspended in deionized (DI) water and deposited overnight onto poly-L-Lysine coated glass coverslips. Next, the collagen-coated substrates are functionalized by covalently attaching the PSMA-encapsulated QDs through EDC-NHS mediated crosslinking. Atomic force microscope (AFM) topographical images of bare collagen nanofibers and hybrid collagen-QD matrices are shown in Figure 2.

**Figure 2.**
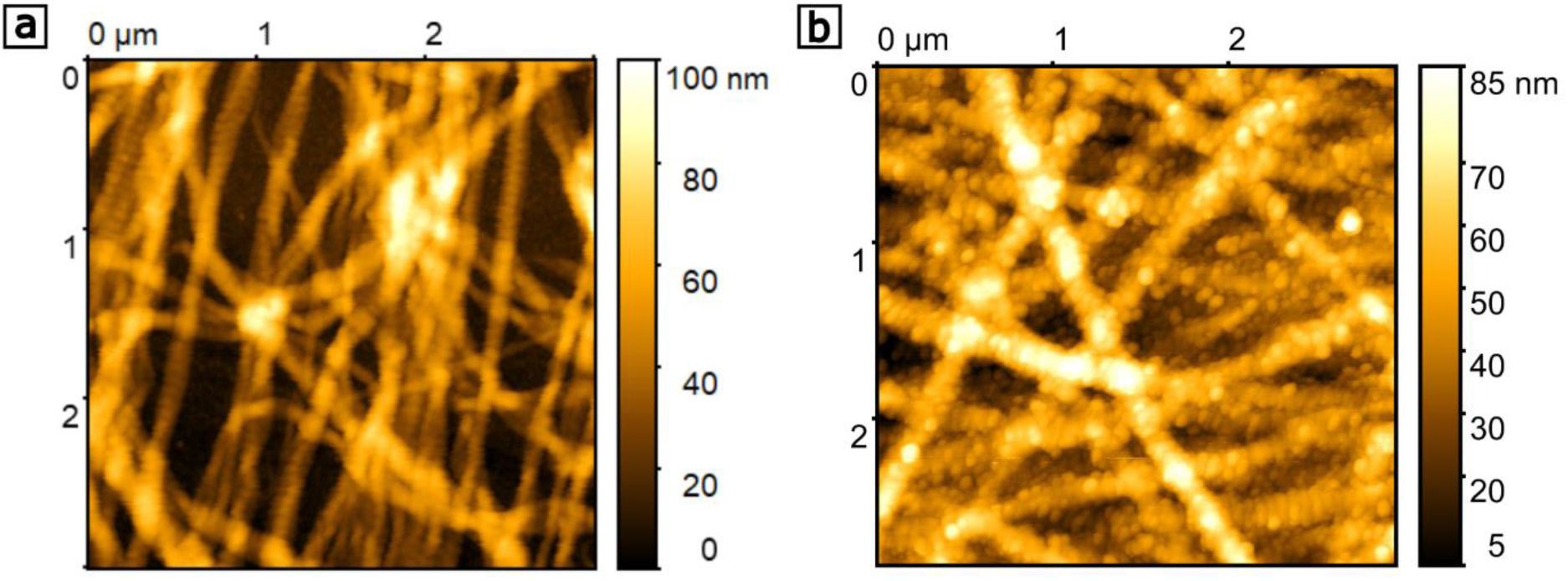
Atomic force microscope (AFM) topographical images of (a) a bare, surface deposited collagen scaffold and (b) a hybrid QD-collagen scaffold.

Finally, the substrates are sterilized using 100% ethanol prior to the culture of rat primary ventricular cardiomyocytes.

### 2.2. Cellular activity monitoring

To validate the optical recordings, a dedicated patch clamp-multiphoton microscopy setup is used to simultaneously measure the cellular electrical activity under whole-cell current-clamp conditions and the two-photon fluorescence (2PF) of the QD-containing matrix. Briefly, the setup consists of a tunable Spectra Physics Insight DS+ femtosecond pulsed laser with 120 fs pulses at an 80 MHz repetition rate. The output power is modulated by a combination of a polarizer and a half-wave plate. The laser is tuned to 900 nm and combined with an Olympus Fluoview FV1200MPE laser scanning microscope. The 2PF is measured in reflection using a 470 nm long pass filter and acquired using Olympus FV10MP photomultiplier tubes (PMTs). The patch clamp rig consists of an Axopath 200B amplifier, an analog to digital converter (DAC, Digidata 1550) and a PS-7000C PatchStar Micromanipulator (Molecular Devices, UK).

A typical experiment is depicted in Figure 3. After 3 days in culture, one regularly beating cardiomyocyte is targeted using a freshly pulled 3-5 MΩ short-tapered, fire-polished borosilicate patch clamp recording pipette and whole-cell current clamp is established to monitor spontaneous electrical activity. Using an excitation laser power of 5-10 mW, an area of ∼ 300 x 300 μm^2^ is then scanned to locate the patched cell and the tip of the patch pipette. A meandering scan line is user-defined in the horizontal plane across the entire body of the target cell. Typically, the length of the scanned line is about 200 μm. Triggered by the patch clamp controller, the 2PF of the QDs is recorded along the defined scanning line at a resolution of 512 x 512 pixels, with a minimal pixel dwell time of 10 μs/pixel and a sampling rate of 150 Hz, while simultaneously recording the electrical activity of the target cardiomyocyte. The scanline in the experiment depicted in Figure 3 encompasses several regions that display changes in optical signal intensity concurrent with the action potentials as measured in current-clamp mode. The areas containing pixels displaying changes in 2PF intensity are indicated in white in panel 3c. For these three labeled regions, 2D images of the scanline versus time are shown in panel 3d. The spatial dimensions (vertical axis) of these scanlines range from 4.1 to 5.9 µm and fluorescence changes occur in both positive and negative direction with respect to the fluorescence at rest. Regions 1 and 3 display decreases in QD 2PF while in region 2, the intensity increases upon membrane depolarization.

**Figure 3.**
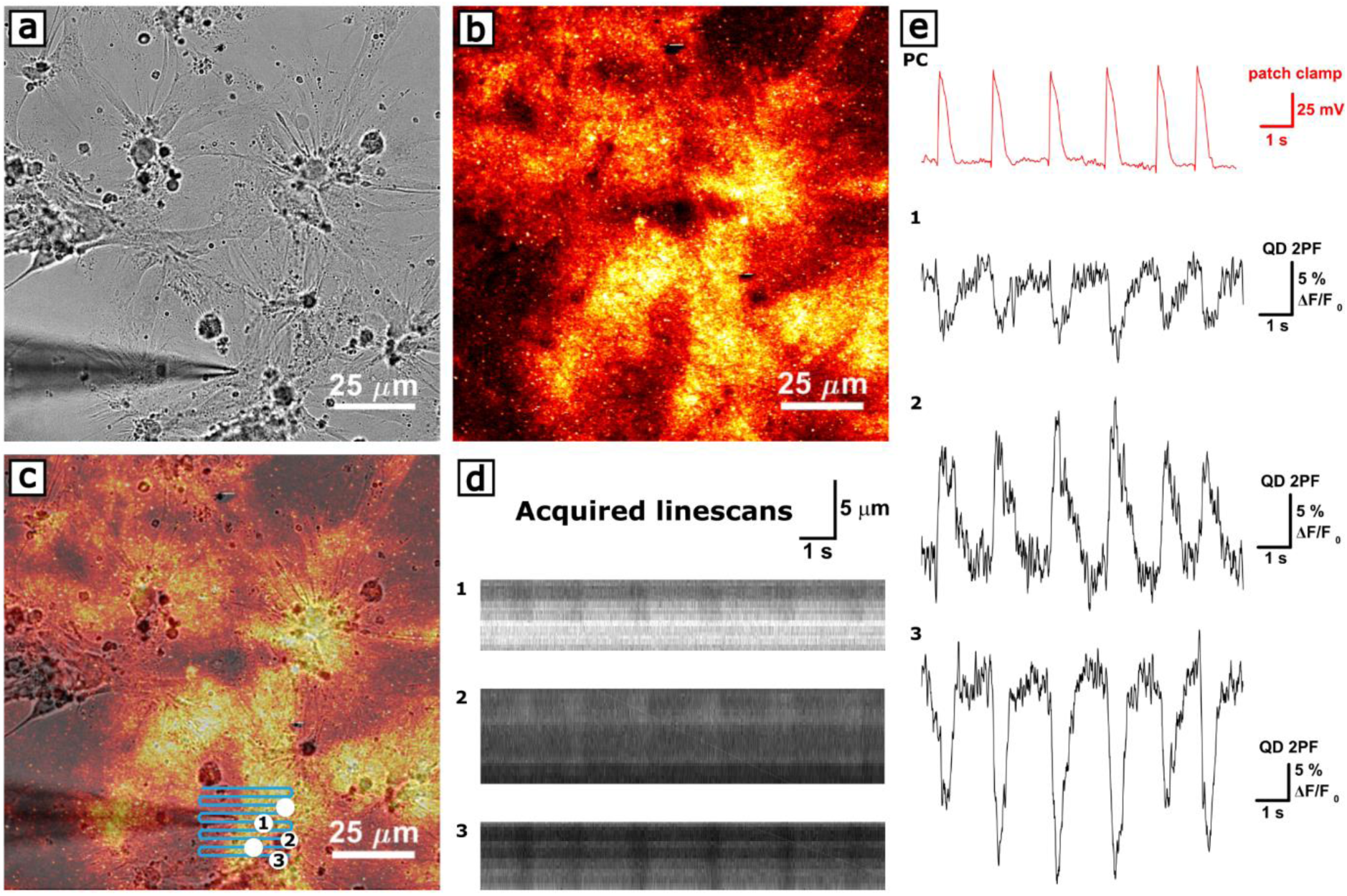
(a) Primary cardiomyocytes cultured (3 days in vitro (DIV)) on top of a QD-functionalized collagen network. One cell is patched in whole-cell current-clamp mode to record spontaneous electrical activity; (b) 2 photon fluorescence (2PF) image of the collagen-quantum dot (QD) layer; (c) Overlay of the bright field microscopy and 2PF images from panel a and b, highlighting the meandering scanline (user-defined) over which the QD 2PF is read out. Regions displaying optical responses are indicated in white; (d) Line scans in the three enumerated regions in panel c are represented as 2D gray scale intensity images where the horizontal axis indicates time and the vertical axis the spatial dimension; (e) QD 2PF traces (black traces) obtained by spatial averaging of the scanlines shown in panel d and the corresponding membrane potential (red trace) as measured in current-clamp mode.

Averaging over the spatial dimension and smoothing with a high-frequency 3^rd^ order Savitzky-Golay filter over an interval of 50 ms, the normalized relative fluorescence change, Δ*F*/*F*_0_ (where Δ*F* = *F* − *F*_0_, *F* representing the time dependent 2PF and *F*_0_ the 2PF at resting potential), is plotted in panel 4e as a function of time together with the membrane voltage traces recorded by the patch clamp electrode.

The cardiomyocyte under investigation in Figure 3 exhibits a stable beat rate of 36 ± 2 bpm as shown by both QD 2PF and electrical recordings. For a membrane voltage change of 75 mV (- 52 ± 4 mV to 23 ± 2 mV), the corresponding relative changes in the QD 2PF responses vary between the different regions, amounting to - 7.3 ± 0.4 %, + 12.8 ± 0.7 % and - 19.5 ± 0.6 % in regions 1, 2 and 3 respectively (averaged over 40 contraction cycles).

A single action potential and the corresponding QD 2PF trace of a regularly beating cardiomyocyte are displayed in more detail in Figure 4. This particular 2PF trace corresponds to a decrease in 2PF intensity as is the case in 62% of all recorded optical responses (of a total of 57 recordings on 23 cells). The relative change in 2PF is location and cell dependent with the top 50% of measurements recording relative changes ranging from 14.0% to 27.1%. Due to this large spread, the mean relative change is not significantly different for positive and negative 2PF variations (p = .067); the relative amplitude for positive changes amounting to 27.52% with a standard deviation of 15.56% (21 recordings) while for negative changes the mean relative change amounts to 20.17% with a standard deviation of 11.22% (36 recordings). In the high signal measurements, the obtained sensitivity is thus comparable to those of newly developed voltage sensitive dyes under two-photon illumination, which can exhibit variations of up to 30 % per 100 mV^31^, or genetic voltage indicators (GEVIs), such as the voltage indicators from the ASAP family (sensitivities up to 25 % in HEK293 under 2P illumination).^1,5,32^ Among the best performing GEVIs under 1P illumination, ASAP2s has responses of up to - 45.1 ± 1.5 % to action potentials in human embryonic derived cardiomyocytes and - 29.1 ± 2.1 % in cardiomyocytes derived from induced pluripotent stem cells^[41]^. In terms of temporal behavior, the full width at half maximum (TD50) for activity recorded in QD 2PF amounts to 432 ± 16 ms as compared to 389 ± 1 ms for an action potential recorded using the patch clamp technique and is independent of the sign and magnitude of Δ*F*/*F*_0_ as shown in Figure 4b.

**Figure 4.**
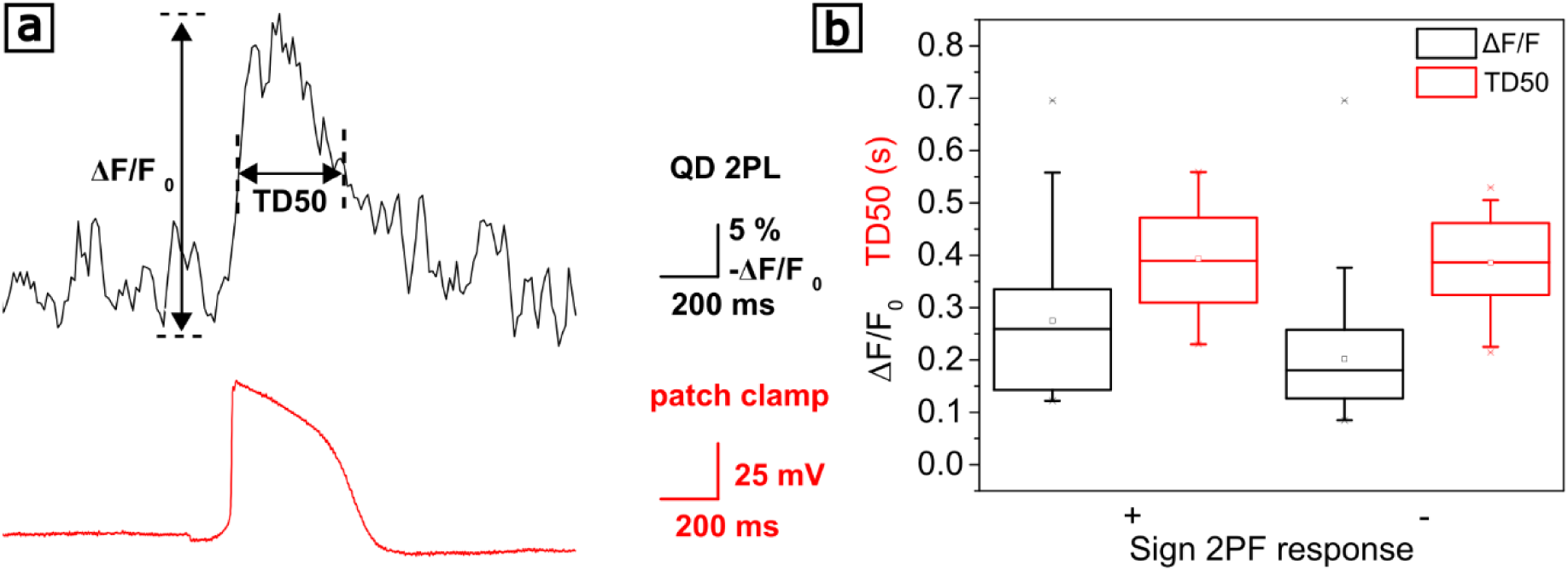
(a) Single action potential (red trace) and corresponding QD 2 photon fluorescence (black trace) at expanded time scale. (b) Maximal relative change and full width at half maximum (TD50) of 2PF cardiomyocyte activity signal for both positive and negative changes.

A direct modulation of the fluorescence of CdSe/ZnS QDs through for instance the Quantum-confined Stark effect (QCSE) is known to result in changes of 1-2% for electric fields up to 150 kV/cm, which is in the order of magnitude of electric-fields over the cell membrane^[42]^. We have attempted to decouple electric-field fluorescence dependence from contraction effects by performing control experiments using the negative inotropic compound blebbistatin. Blebbistatin inhibits actin-myosin interaction and thus suppresses contraction and without influencing electrophysiology^[43]^. When 10 µM blebbistatin is added to the extracellular medium, cell contraction is completely suppressed together with all fluorescence changes (see Figure S2). This experiment would support the hypothesis that the measured fluorescence changes are solely produced by the mechanical deformation of the matrix through the contractile motion of the cardiomyocytes. Nevertheless, we cannot exclude electric field effects since blebbistatin modifies the cell membrane tension and focal adhesions^[44–46]^. It could also modify the QD localization with respect to the cell membrane. Further experiments on non-contracting cells have to be performed in order to conclusively deconvolute these effects.

### 2.2. Evaluation of the effect of epinephrine on contractile parameters

To assess the performance of the developed hybrid collagen - QD scaffolds in monitoring cardiac contractility parameters during drug screening, we studied the effect of epinephrine, which has a positive chronotropic, inotropic as well as lusitropic effect, increasing respectively the beat rate, contractile force and rate of myocardial relaxation. Changes in 2PF signals for different epinephrine concentrations are shown in Figure 5 together with patch clamp recorded data.

**Figure 5.**
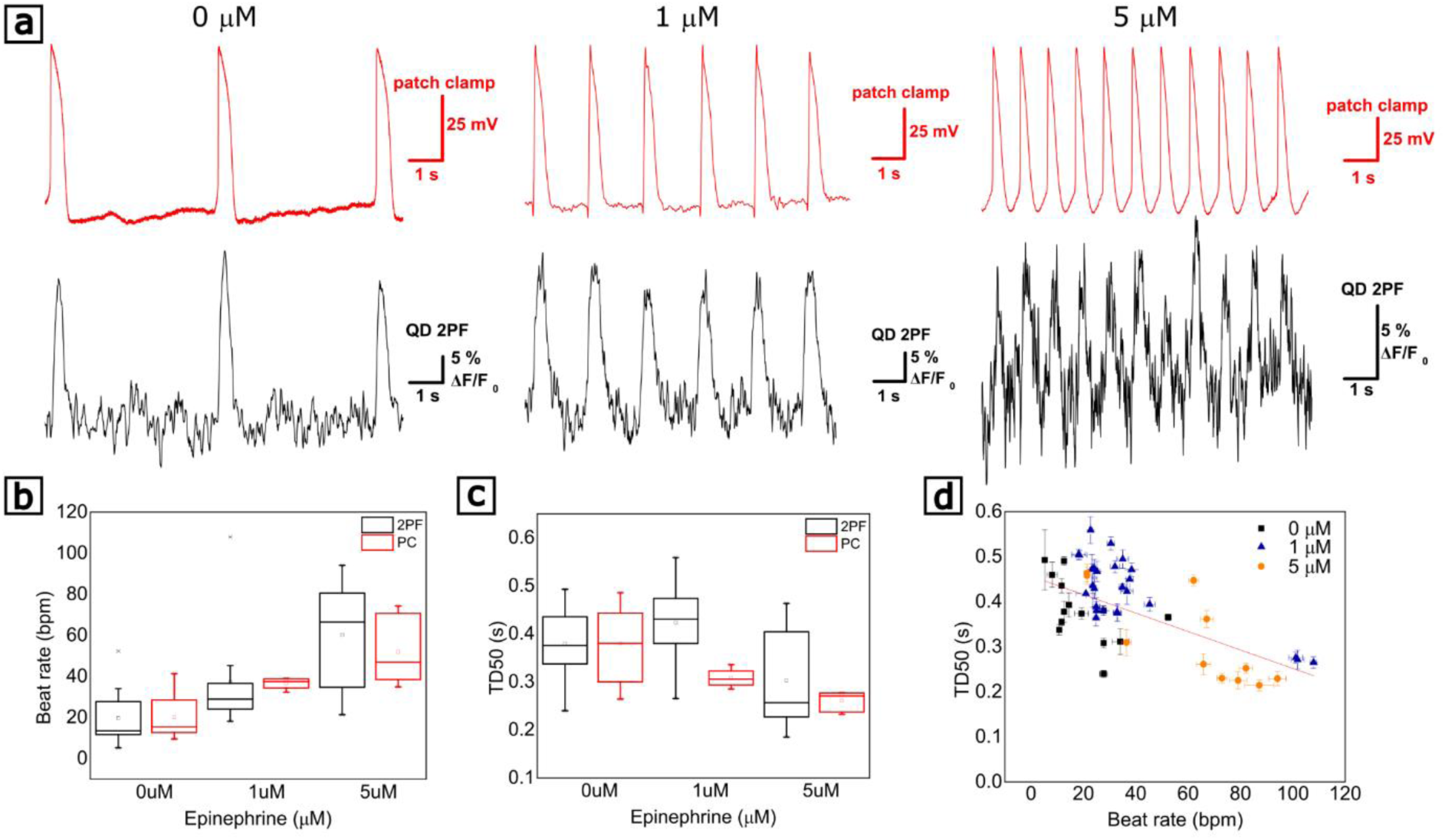
Drug-induced effects on cardiac contractility parameters measured by QD 2PF. (a) Signals measured in the absence of epinephrine and increase in the cardiomyocyte beat rate in the presence of 1 µM and 5 µM epinephrine. Electrical recordings (red traces) and corresponding 2 photon fluorescence (2PF) recordings (black traces). (b) Beat rate in beats per minute (bpm) versus epinephrine concentration in µM as measured by current clamp and QD 2PF in independent measurements; (c) Half width at full maximum (TD50) in seconds for current clamp and QD 2PF measurements; and (d) TD50 in seconds versus beat rate in bpm grouped for different concentrations of epinephrine (0 µM, 1 µM and 5 µM). The fit has a slope of -2.1 ± 0.4 ms / bpm.

Chronotropically, upon exposure to concentrations of 1 µM and 5 µM epinephrine, the beat rate increases from 19.7 ± 3.5 bpm to 37.3 ± 5.0 bpm at 1 µM and 62.7 ± 7.7 bpm at 5 µM, respectively, in excellent agreement with the electrical activity as measured in current clamp. Traces of the electrical activity as measured by current clamp and QD 2PF for the three distinct scenarios are shown in Figure 5a. Secondly, the lusitropic effect of epinephrine causes a measurable change in TD50 with values ranging of 0.42 ± 0.02 s and 0.31 ± 0.03 s, for concentrations of 1 µM and 5 µM respectively. Figure 5d clearly shows that the positive lusitropic effect correlates with beat rate. The TD50 value decreases linearly with 21 ± 4 ms per increase of 10 bpm in beat rate. A large spread in recorded amplitudes, as discussed earlier in Figure 4, prohibits the quantification of inotropic effects. In addition to epinephrine, this is also tested using the negative inotropic agent blebbistatin.

Nevertheless, this demonstrates that through recording of the 2PF of the QDs, cardiomyocyte beat rate and contractile relaxation can be monitored and quantified.

## 3. Conclusions and perspectives

This study demonstrates the possibility of reading out cardiomyocyte activity through changes in 2PF of a QD-decorated fibrous collagen ECM. The optical signal reflects the mechanical deformation of the matrix through contractile motion of the cardiomyocytes and it is consistent with electrical recordings. The approach is validated as a tool to detect and quantify epinephrine-induced chronotropy and lusitropy via cardiac contractility parameters. Control experiments using blebbistatin to block contractile activity did not indicate the presence of a direct electric field modulation of the 2PF signal through for instance the Quantum-confined Stark effect. However, as mechanical and electrical activity cannot be easily uncoupled in the case of cardiomyocytes without affecting other important parameters (such as the membrane tension), further studies should be performed on non-contracting electrogenic cells – such as neurons. Neural activity recording imposes much more stringent requirements on the time resolution (below millisecond), which are not attainable with our current set-up. Improved time resolution should be attainable by controlling the dimensions of the scan line and moving towards single voxel read-out. Moreover, random-access multiphoton microscopy would allow one to measure 2PF changes at several distinct locations by fast beam movement^[41,47]^. Finally, although we only prepared 2D primary rat cardiomyocyte cell cultures, the concept should be easily applicable to 3D tissue constructs and should therefore be suitable for *in situ*, real-time monitoring of electrical activity in developing cardiac tissues.

## 4. Materials and Methods

All chemicals were obtained from Sigma Aldrich unless stated otherwise.

### QD phase transfer

In a typical phase transfer experiment, the excess ligands of a 10 μM octadecylamine/trioctylphosphine (TOPO) stabilized CdSe/ZnS core-shell QD solution (Sigma Aldrich product no 790206) in chloroform (CHCl3) were removed by a twofold centrifugation of an equal volume of QD solution and a 1:1 methanol:aceton mixture (Acros Organics). Next, 20 mg of poly-styrene-co-maleic anhydride (PSMA) were mixed with 200 μL of the QD solution. After 4 hours of gentle shaking, 0.1 mL of ethanolamine in water was added. The ethanolamine opened the exposed anhydride rings of the PSMA, thereby producing carboxylic acid groups. Vigorous shaking then transferred all the QDs to the water phase. The water-soluble QDs were collected, purified using a PD 10 desalting column (GE Healthcare) to remove excess PSMA and stored in pH 9, 10 mM borate.

Quantum yield (QY) values were determined using the comparative method described by Würth et al.^[48]^ using Rhodamine 6G as reference dye with a known QY of 0.96 in water^[49]^. Fluorescence and absorbance spectra were measured using a Tecan infinite 200PRO microwell plate reader (Tecan Trading AG, Männedorf, Switzerland).

Dynamic light scattering (DLS) measurements were performed using a NanonBrook 90plus particle size analyzer (Brookhaven, Holtsville, N.Y., USA) at a detection angle of 90°.

### Scaffold preparation

Borosilicate glass coverslips (25mm diameter, VWR) were cleaned for 20min in piranha solution (1:3 v/v H2O2 (30 %) H2SO4 (98 %)), rinsed with ultrapure water and dried under nitrogen flow. Subsequently, the clean coverslips were coated with a 0.1 mg/ml poly-L-lysine (PLL) solution in pH 8.5, 100 mM borate buffer.

3D collagen gels were prepared through the self-assembly of monomeric collagen from bovine skin. by adjusting the pH of the acidic solution of monomeric collagen to 7.4 using phosphate-buffered saline (PBS) and raising the temperature to 37 °C for 4 hours. Next, the fiber network was broken up into single fibers through horn sonication (Branson sonifier 150, 1W) for 10s, centrifuged and resuspended in water. 200 µL of the resulting collagen fiber solution was then deposited overnight on a PLL coated coverslip.

Finally, the water solubilized QDs were covalently bound to the collagen matrices. 100 □L of 0.2 mg/ml PSMA encapsulated QDs was mixed with 100 μL of a 5 mM 1-Ethyl-3-(3-dimethylaminopropyl)-carbodiimide (EDC) and 5 mM sulfo-N-hydroxysulfosuccinimide (NHS) mixture in pH 6.5, 0.1 M MES buffer. The mixture was allowed to react for 30 min and then 200 μL were applied to a collagen coated coverslip. After 2 h incubation, unreacted QDs were rinsed off and the sample was dried under nitrogen flow.

Atomic force microscopy (AFM) images of collagen-QD scaffolds are obtained using a JPK instruments bioafm Nanowizard NanoOptics atomic force microscope using PPP-NCHR tapping mode AFM probes (Nanosensors, Neuchâtel, Switzerland).

### Cell culture

Neonatal rat ventricular cardiomyocytes were harvested from euthanized two days old Wistar rats (protocol approved by the KU Leuven animal ethics committee). The ventricles were cut into pieces, washed in Hanks balanced salt solution (HBSS) and incubated overnight in 0.05 % trypsin. Next, the tissue was digested using 1 mg/ml collagenase, followed by mechanical trituration and two centrifugation steps. The cell suspension was first centrifuged at 300g for 5 min, the pellet resuspended in HBSS containing 6 % BSA, then centrifuged at 400g for 4 min and finally the cells were dissociated in culture medium (Ham F10 containing 5 % FCS, 1 % Penicillin-Streptomycin, (4-(2-hydroxyethyl)-1-piperazineethanesulfonic acid (HEPES), 0.5 % ITS, 0.1 mM Norepinephrine, and 2 μg/ml vitamin B-12). After a pre-plating step in a T-75 cell culture flask to remove remaining fibroblasts, the cardiomyocytes were seeded on the collagen-QD substrates at a density of ∼ 40 %.

### Cell electrophysiology

Whole-cell patch clamp recordings from rat cardiomyocytes were performed using 2-5 MΩ borosilicate glass pipettes with filaments (BF150-86-7.5), pulled on a P-1000 Flaming/Brown Micropipette Puller (Sutter Instruments, California, USA). The extracellular recording solution (ECM) contained 1.8 mM CaCl2, 15 mM glucose, 5.4 mM KCl, 1 mM MgCl2, 150 mM NaCl, 1 mM Na-pyruvate, 15 mM HEPES at pH 7.4. Additionally 10 µM (±)- blebbistatin or 3-5 µM (±)-epinephrine hydrochloride was supplemented. The micropipette was filled using nonmetallic syringe needle (MicroFill, World Precision Instruments, Florida, USA) with an intracellular solution (ICM) containing 2 mM CaCl2, 150 mM KCl, 5 mM NaCl, 5 mM egtazic acid (EGTA), 5 mM MgATP and 10 mM HEPES at pH 7.2. Both solutions were sterile filtered with an 0.2 μm syringe filter and had a final osmolarity 310 (intracellular) and 330 (extracellular) mOsm (Osmomat 3000 basic, Gonotec Germany). Cells were images using an Olympus BX61 W1 microscope fitted with a 40x water immersion objective (0.80 NA, 3.5 mm WD, Nikon). The microscope was coupled to a tunable mode-locked femtosecond laser (Insight DS, Spectra Physics) with a repetition rate of 80 MHz and pulse widths of 120 fs, set to an excitation wavelength of 900 nm. The laser output power was set to 5-10 mW by combining a polarizer and achromatic half-wave plate. Filtered through a 450 nm long-pass filter, the two-photon luminescence was measured in reflection using Olympus FV10MP photomultiplier tubes.

The multiphoton microscope was combined with a patch clamp setup consisting of an Axopath 200B amplifier, an analog to digital converter (DAC, Digidata 1550 low nofafise acquisition system) and a PS-7000C PatchStar Micromanipulator Molecular Devices, UK. The recording electrode was a chlorinated silver wire (0.25 mm thickness, 1-HLA-005, Molecular Devices, California, USA).

## Acknowledgements

S. Jooken acknowledges the financial support by the Flanders Research Foundation (FWO) - strategic basic research doctoral grant 1SC3819N. C. Bartic, G. Callewaert and O. Deschaume acknowledge the financial support by the Flanders Research Foundation (FWO grant G0947.17N) and KU Leuven research grants OT/14/084 and C14/18/061. T. Verbiest acknowledges financial support from the Hercules Foundation.

## Supporting Information

**Figure S1.**
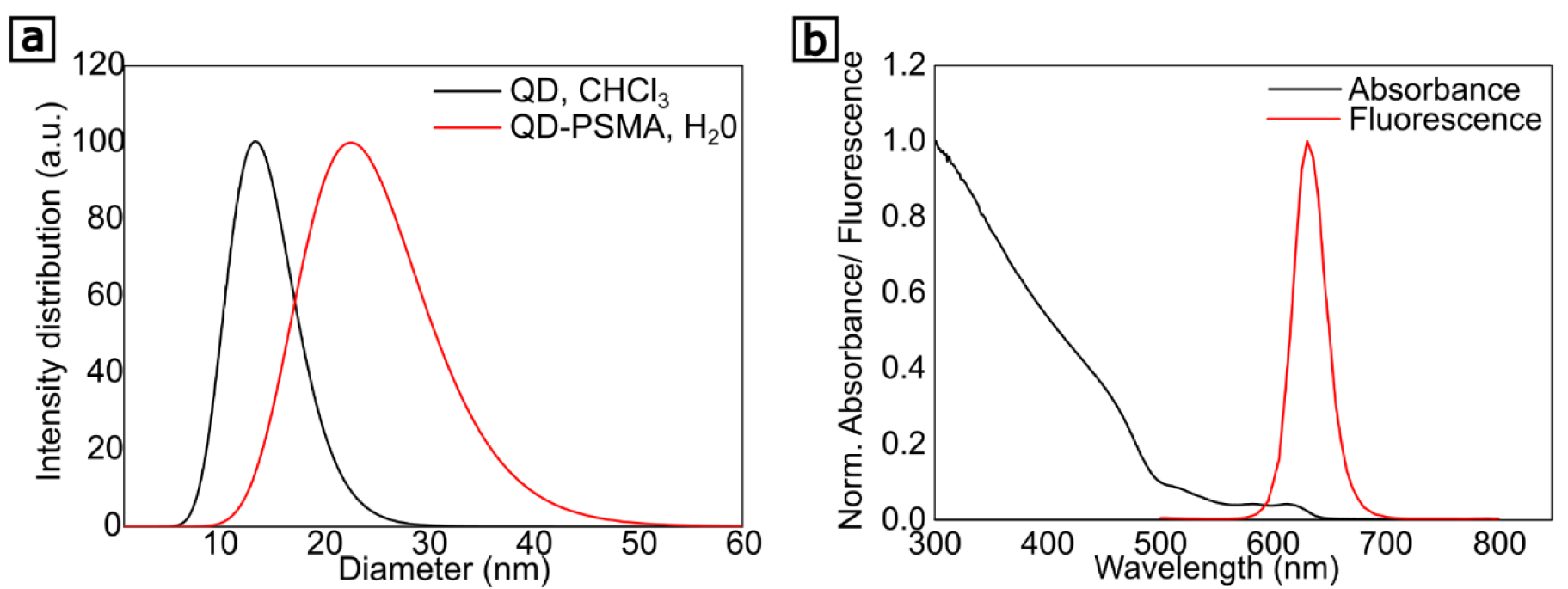
(a) Lognormal size distribution as measured by dynamic light scattering (DLS) of hexadecylamine (native ligands) capped CdSe/ZnS QDs in chloroform (CHCl3) (black curve) and water transferred poly-styrene-co-maleic-anhydride (PSMA) encapsulated QDs. (b) Fluorescence (red) and absorbance spectra (black) of the PSMA encapsulated QDs.

**Figure S2.**
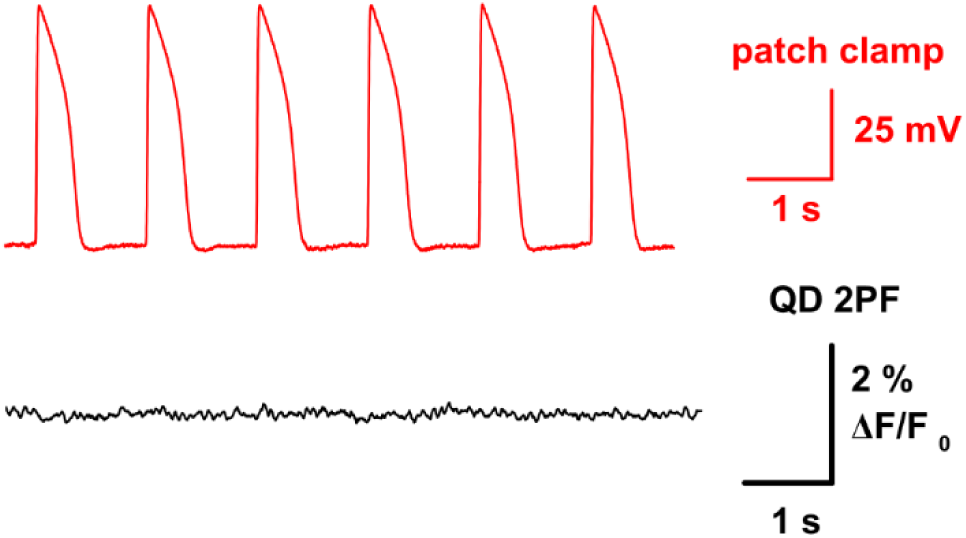
Typical measurement of the membrane potential (red trace) as measured by whole-cell current clamp and QD 2PF read-out (black trace) of a cardiomyocyte with 10 µM blebbistatin supplemented to the extracellular medium.

